# Vision using multiple distinct rod opsins in deep-sea fishes

**DOI:** 10.1101/424895

**Authors:** Zuzana Musilova, Fabio Cortesi, Michael Matschiner, Wayne I. L. Davies, Sara M. Stieb, Fanny de Busserolles, Martin Malmstrøm, Ole K. Tørresen, Jessica K. Mountford, Reinhold Hanel, Kjetill S. Jakobsen, Karen L. Carleton, Sissel Jentoft, Justin Marshall, Walter Salzburger

## Abstract

Vertebrate vision is accomplished through a set of light-sensitive photopigments, which are located in the photoreceptors of the retina and consist of a visual opsin protein bound to a chromophore. In dim-light, vertebrates generally rely upon a single rod opsin (RH1) for obtaining visual information. By inspecting 101 fish genomes, we found that three deep-sea teleost lineages have independently expanded their *RH1* gene repertoires. Amongst these, the silver spinyfin (*Diretmus argenteus* Johnson 1863) stands out as having the highest number of visual opsins known for animals to date (2 cone and 38 rod opsins). Spinyfins simultaneously express up to 14 *RH1s* encoding for photopigments with different peak spectral sensitivities (λ_max_=448-513 nm) that cover the range of the residual daylight, as well as the bioluminescence spectrum present in the deep-sea. Our findings present novel molecular and functional evidence for the recurrent evolution of multiple rod opsin-based vision in vertebrates.

**SHORT ABSTRACT:** Contrary to the single rod opsin used by most vertebrates, some fishes use multiple rod opsins for vision in the dimly lit deep-sea.

---

Animals use vision for a variety of fundamental tasks including navigation, food acquisition, predator avoidance, and mate choice. At the molecular level, the process of vision is initiated through a light-induced conformational change in a photopigment (a visual opsin protein bound to a vitamin A-derived chromophore)^1-3^, which in turn activates the phototransduction cascade^3,4,5^. Vertebrates possess up to five basic types of visual opsins, of which four are primarily expressed in cone photoreceptors in the retina (‘cone opsins’), whereas a fifth one is expressed in the rod photoreceptor (‘rod opsin’)^2,6,7^. Cones generally operate in bright-light (photopic) conditions and are sensitive to a broad range of wavelengths^7,8^: Photopigments containing the short-wavelength-sensitive opsins, SWS1 and SWS2, absorb in the ultraviolet (UV, λ_max_=355-450 nm) and violet/blue (λ_max_=415-490 nm) regions of the light spectrum, respectively; the middle-wavelength-sensitive pigment (RH2) is most sensitive to the central (green) waveband (λ_max_=470-535 nm), whereas the long-wavelength-sensitive pigment (LWS) is tuned towards the red end of the spectrum (λ_max_=490-570 nm)^2,8,9^. Usually, vertebrates rely on two to four spectrally distinct cone photoreceptors for colour opponency, that is, the ability to distinguish different chromatic signals^1^. Under dim-light (scotopic) conditions, however, most vertebrates are colour-blind, relying upon their single rod photopigment (RH1) for obtaining achromatic visual information^10,11^.

In this study, we scrutinised the evolution of the visual opsin-gene repertoire of teleost fishes, with a particular focus on deep-sea fishes inhabiting the mesopelagic zone below a depth of 200 m. Deep-sea fishes exhibit various adaptations to maximise their visual sensitivities in a scotopic environment where bioluminescence replaces surface illumination as the primary source of light^12,13^. Among these adaptations are changes in eye morphology, such as an increase in eye size, pupil size, reflective tapeta or extremely modified tubular eye structures^14–17^. Other modifications include changes to the retina itself^18^. For example, it has been known for some time that the retina of the silver spinyfin (*Diretmus argenteus*) is composed of morphologically distinct rod photoreceptors stacked in multiple layers^19^. While the exact function of this ‘multibank retina’ is unknown, it was previously speculated that the different layers could be used to replenish deteriorating rods from subjacent layers or to discriminate colour in the deep sea via retinal tiering^20^.

To examine molecular adaptations in the visual system of teleosts, we first reconstructed and, in some instances, re-assembled the visual opsin gene loci in 100 teleost genomes (plus one non-teleost outgroup; Fig 1; Fig. S1; Table S1). We found that teleosts possess a median number of seven visual opsin genes per species. The elevated number of visual opsins in teleosts compared to other vertebrates^5^ can primarily be attributed to an expansion of *SWS2* and *RH2*, which form photopigments that maximally absorb in the most frequent, blue-green part of the aquatic light spectrum. Specifically, we found that 78 of the 100 teleost species examined had more than one copy of *RH2*, and 53 species had at least one extra version of *SWS2* (see also ref. 21) (Fig. 1; Fig. S1). Gene losses, on the other hand, mainly affected the cone opsins sensitive to the edges of the visible light spectrum, *SWS1* (absent in 46 species) and *LWS* (absent in 34 species, of which 28 live in the deep sea) (Fig. 1; Fig. S1). A phylogenetic generalised least squares (PGLS) analysis revealed that, while the total number of visual opsin genes was unaffected by phylogeny or the depth at which a species lives (PGLS; Pagel’s λ=0, F_1,74_=0.32, p=0.57), deeper-dwelling species have significantly fewer *LWS* genes (PGLS; Pagel’s λ=1.0, F_1,74_=38.47, p<0.0001) (Table S2).

**Figure 1.**
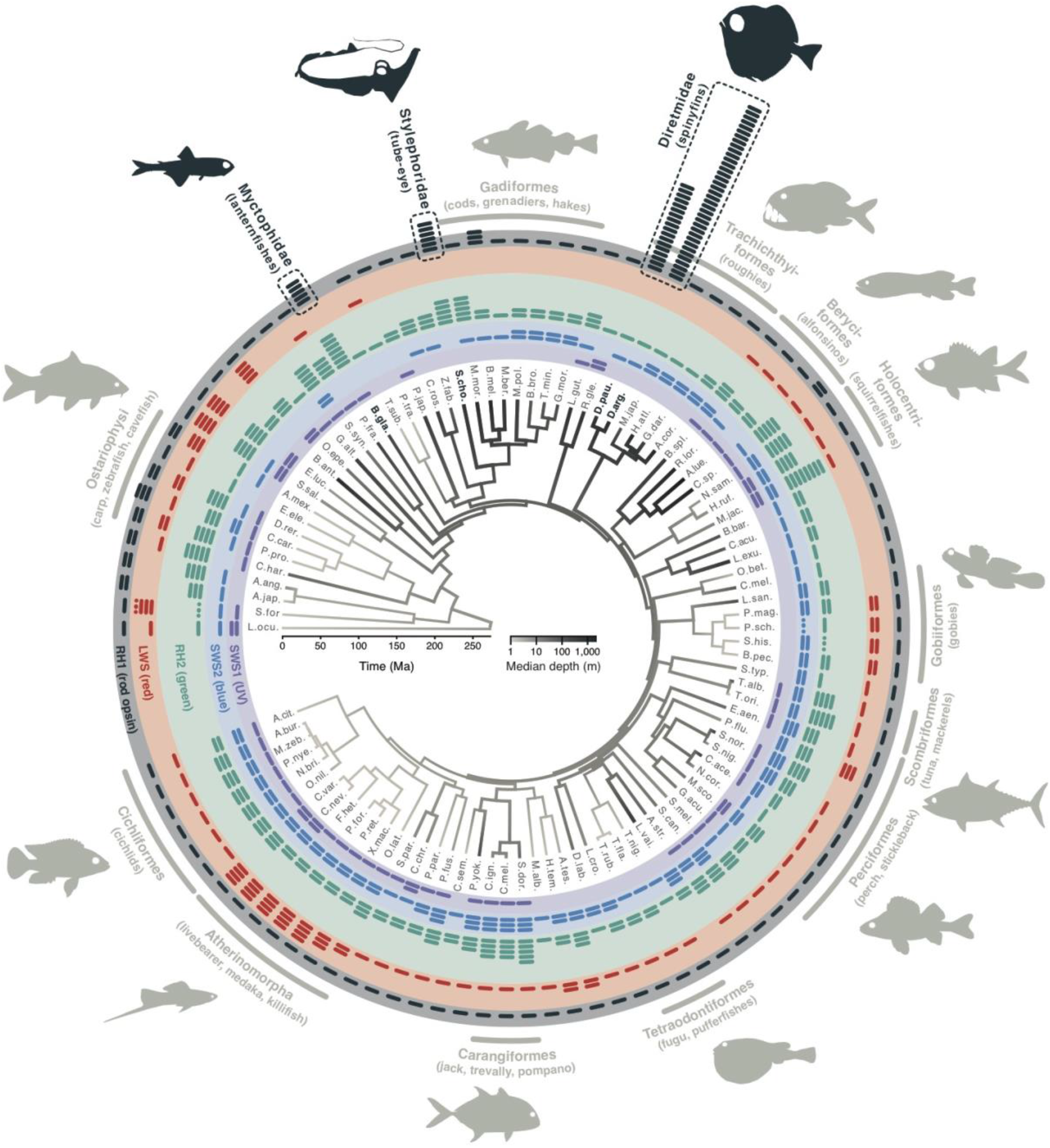
Diversity of visual opsin genes in teleost fishes. The time-calibrated phylogeny in the centre is based on molecular information provided by 101 fish genomes and shaded according to the median depth of occurrence of each species (terminal branches) and reconstructed depths (internal branches). The coloured bars in the outer circles indicate the number of cone opsin genes in the respective genome (whereby *SWS1* is coloured in violet, *SWS2* in blue, *RH2* in green, and *LWS* in red, in line with the genes’ wavelength of maximal light absorbance), while the number of rod opsin genes (*RH1*) is shown as black bars; dotted bars refer to incomplete or ambiguous data. Deep-sea lineages with multiple *RH1* gene copies are highlighted with dashed boxes, amongst which the silver spinyfin (*Diretmus argenteus*) stands out as having the highest number of visual opsins known in animals. A detailed version of the phylogeny including full species names and information on phylogenetic inference, gene synteny, and orientation of the genes in the genome is provided in Fig. S1.

The genome data also revealed, that like most vertebrates^5^, the majority of teleosts possess a single rod opsin gene (Fig. 1) irrespective of phylogeny and depth (PGLS; Pagel’s λ=0, F_1,74_=2.87, p=0.09) (Table S2). On the other hand, we identified 13 species with more than one *RH1* gene (Fig. 1; Fig. S1). Out of these, four deep-sea species from three phylogenetically distinct lineages stand out by possessing five or more *RH1* genes: The glacier lanternfish (*Benthosema glaciale*; Myctophiformes) has 5 *RH1* genes and the tube-eye (*Stylephorus chordatus*; Stylephoriformes) has 6 *RH1*s, which is outnumbered by the *RH1* gene repertoire of two species of the family Diretmidae (Trachichthyiformes), the longwing spinyfin (*Diretmoides pauciradiatus*) with 18 functional *RH1* genes, and the silver spinyfin (*Diretmus argenteus*) with 38 *RH1*s (Fig. 1). Interestingly, in all cases, the *RH1* gene expansions occurred through repeated single gene- rather than whole-genome duplications (Fig. S2; Table S3).

To determine how many and which of the visual opsin genes are actually being used, we sequenced the retinal transcriptomes of 36 species distributed across the phylogenetic spectrum of teleosts (Table S1). We found that the majority of species (n=24) express up to four cone opsins, despite having – in many cases – more than four cone opsin genes in the genome (Fig. S3). This is consistent with the use of two to four differently tuned cone photoreceptors for colour vision^5,22^ on land and in freshwater environments^23^. On the other hand, the retinae of the remaining 12 species contained transcripts of five to seven cone opsin genes, potentially challenging this view (Fig. S3). Regarding the rod opsin, we found that deep-sea fishes with an extended *RH1* repertoire indeed express more than one *RH1* gene in their retinal tissue. For example, the two lanternfishes *B. glaciale* and *Ceratoscopelus warmingii* express three *RH1* genes each, and the tube-eye (*S. chordatus*) expresses five *RH1s* in its retina (Fig. S3). Moreover, on top of expressing one or two distinct cone opsins, the silver spinyfin (*D. argenteus*) was found to express up to seven and fourteen functional rod opsins as larvae and adults, respectively (Fig. S3), with four of the *RH1* transcripts found in both the developmental stages (Fig. 2a). A similarly high number of opsin transcripts has previously only been reported from dragonflies^24^ and stomatopod crustaceans^25^, in the latter case generating up to 12 differently tuned photoreceptor types for colour vision.

**Figure 2.**
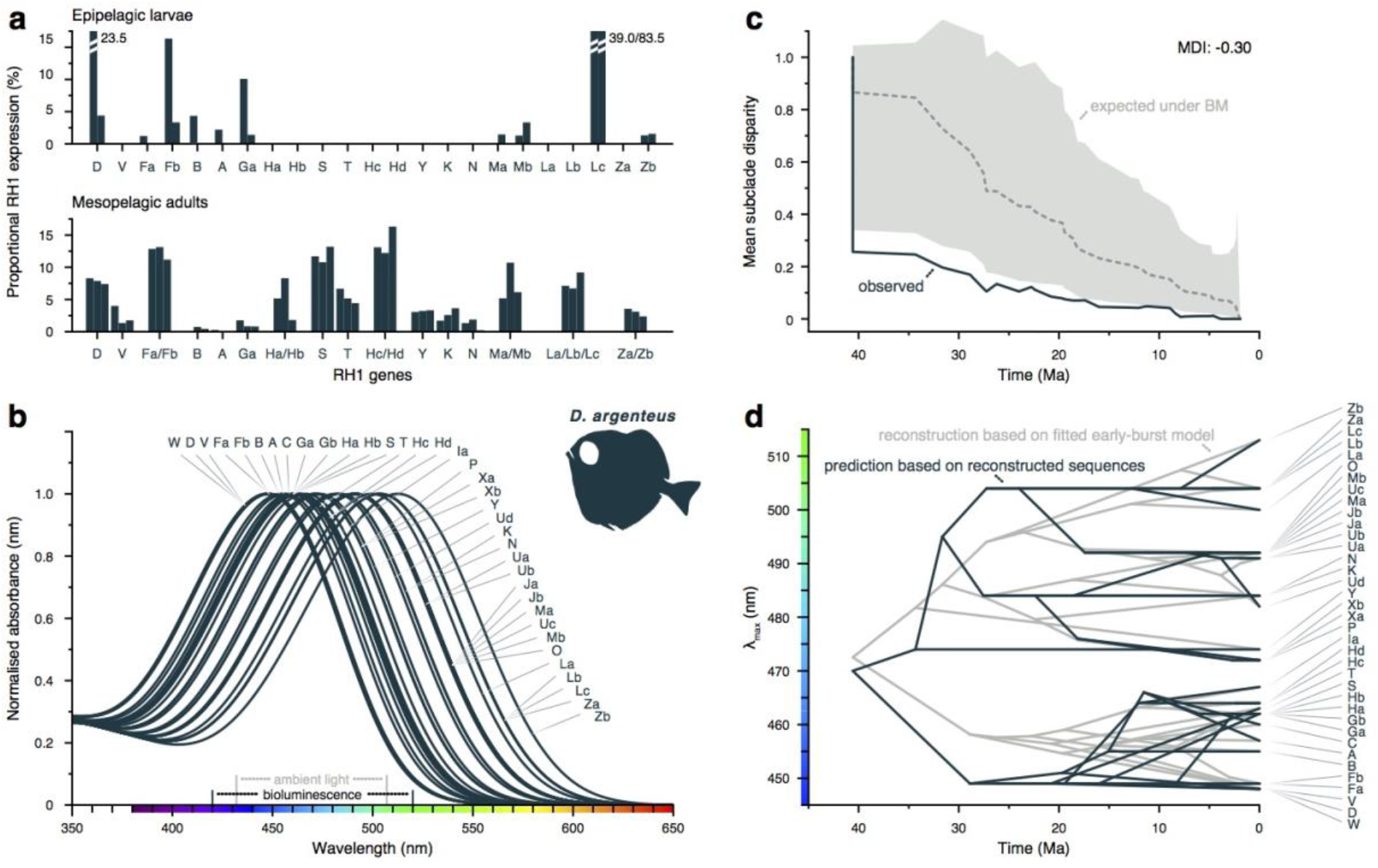
Molecular function of rod photoreceptors in the silver spinyfin (*Diretmus argenteus*). **a**, Expression of rod opsin (*RH1*) genes in the retina of epipelagic larvae (upper panel) and adults (lower panel) of *D. argenteus*. Note, that no association was found between gene expression and differences in transcription factor binding sites upstream of the *RH1* genes (Table S8). **b**, Peak spectral sensitivities (λ_max_) of 37 (out of the 38) RH1 photopigments of *D. argenteus* based on *in vitro* protein regenerations and subsequent functional predictions. Note that the λ_max_-values of *D. argenteus* RH1s cover the waveband of deep-sea bioluminescence^13^ and residual daylight present at a depth of 500 m^29^. **c**, Reconstruction of mean sub-clade disparity in λ_max_ through time for the RH1 photopigments of *D. argenteus* supporting an early burst (EB) scenario of diversification in λ_max_ (AIC_EB_=236.4) over time-homogenous diversification (Brownian Motion, BM; AIC_BM_=244.9) or selection towards an optimal value of λ_max_ (Ornstein-Uhlenbeck model, OU, not shown; AIC_OU_=246.9). **d**, Functional divergence of λ_max_ through time according to predicted λ_max_ for reconstructed ancestral sequences (black) and ancestral λ_max_ reconstructed on the basis of the early-burst model with the best fit to the current λ_max_ values (grey). MDI: morphological disparity index.

In vertebrates, substitutions at 27 amino acid positions have so far been implicated with the spectral tuning of RH1 photopigments via functional shifts in λ_max_ (ref. 8 and Supplementary Material & Methods). Our ancestral state reconstruction revealed that 25 out of these 27 known key spectral tuning sites have been altered across the teleost phylogeny (Fig. 3a, c; Fig. S4), and at 18 of these key spectral tuning sites, the same amino acid substitutions have occurred repeatedly and in parallel in different teleost lineages (Table S4). Interestingly, the lineage-specific expansion of *RH1* within Diretmidae alone gave rise to a set of genes differing in 24 out of the 27 key spectral tuning sites (Tables S4, S5). In addition, the *RH1* genes of Diretmidae show the, by far, highest non-synonymous to synonymous (dN/dS) substitution rates across all teleost *RH1*s (Fig. 3b; Table S6), suggesting that the extensive occurrence of parallel substitutions at the molecular level was driven by adaptive sequence evolution.

**Figure 3.**
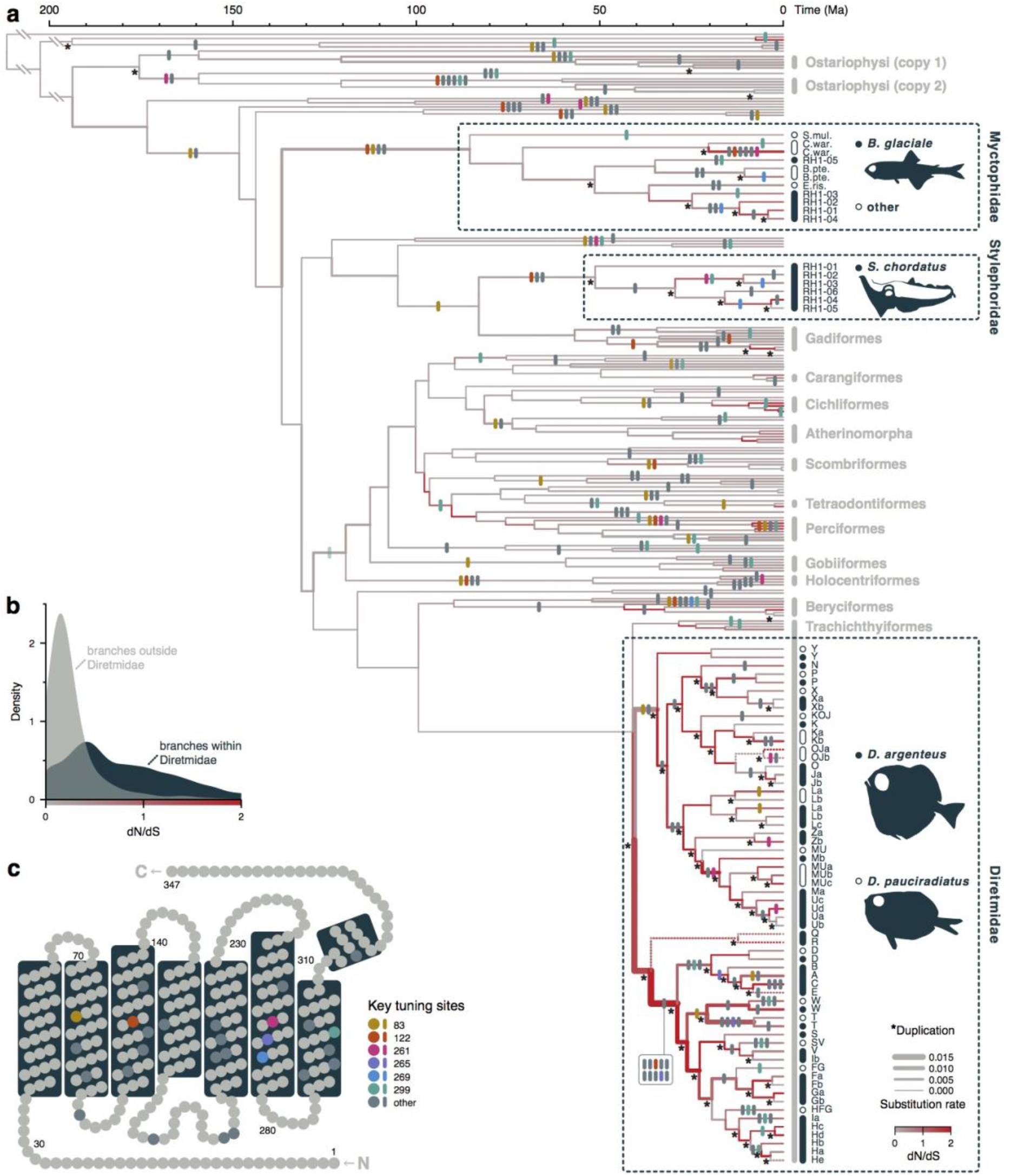
Evolution and functional diversification of RH1 in deep-sea fishes. **a**, Time-calibrated gene tree based on teleost rod opsins (*RH1*) demonstrating lineage-specific rod opsin duplications in three deep-sea fish lineages. Vertical bars indicate amino acid substitutions in key spectral tuning sites^2,5,10,24^ and following the sites highlighted in **c**. Gene duplication events are marked with an asterisk. The branches in the gene tree are colour-coded according to the rate of non-synonymous to synonymous substitutions (dN/dS); the thickness of each branch refers to the reconstructed substitution rate. **b**, Distribution of the per-branch dN/dS-values within the *RH1* genes of Diretmidae compared to all other branches in the teleost *RH1* gene tree based on ancestral sequence reconstructions **c,** Basic model of the RH1 protein showing its seven trans-membrane helices and the positions of the known key spectral tuning sites.

To test whether the exceptional diversity in *RH1* in *D. argenteus* translates into a range of λ_max_-values, we regenerated a subset of *D. argenteus* RH1 photopigments *in vitro* (Fig. S5). Measurements of λ_max_ of six proteins and subsequent predictions for an additional 31 *RH1* genes showed that the *D. argenteus* rod opsin visual pigments cover a range of 448 nm to 513 nm in λ_max_ (Fig. 2b, Table S7). This peak-to-peak spectral range of 65 nm is much broader than that commonly found for RH1 in other deep-sea fishes, in which λ_max_-values range from 477 nm to 490 nm^26–28^. However, this spectral range largely overlaps with both the waveband of bioluminescent light emitted by deep-sea organisms (λ=420-520 nm)^13^ and the residual daylight at a depth of 500 m (λ=432-507 nm)^29^. Interestingly, our function-through-time analysis revealed that the ancestral *RH1* gene of Diretmidae encoded for a photopigment with a λ_max_-value of ∼472 nm, which is similar to the spectral peak of the rod photopigment in most extant deep-sea fishes^27^ (Fig. 2c, d). A subsequent gene duplication event then led to two versions of *RH1* that diverged rapidly towards substantially different spectral sensitivities (λ_max_∼457 nm and 482 nm). These ancestral versions gave rise to all further *RH1* genes that now cover the full extent of the ambient-light spectrum in the deep sea (Fig. 2d).

Two different but not mutually exclusive scenarios could explain the expansion of RH1 photopigments in deep-sea fishes, both of which have not been reported in vertebrates to date. First, similar to nocturnal invertebrates such as flour beetles^30^, deep-sea fishes may coexpress multiple differently tuned RH1 photopigments within the same photoreceptor to boost sensitivity across the ambient light spectrum. So far, however, vertebrates have been found to coexpress a maximum of three (cone) opsin genes per cell^31^, a small fraction of the number of opsin genes expressed by *D. argenteus*. The vastly expanded opsin-gene repertoire of Diretmidae, and *D. argenteus* in particular, is therefore intriguing, especially in the context of its retinal anatomy: the ventral part of the *D. argenteus* retina contains extremely long rods as part of a multi-bank retina (Fig. 4a–c)^19,20^. Placing just the shortest and the longest tuned *D. argenteus* visual pigment within these rods results in very broad absorptance spectra that would not only overlay the spectra of the remaining RH1 pigments, but would already on its own maximise photon capture across the deep-sea light spectrum (Fig. 4d, e).

**Figure 4.**
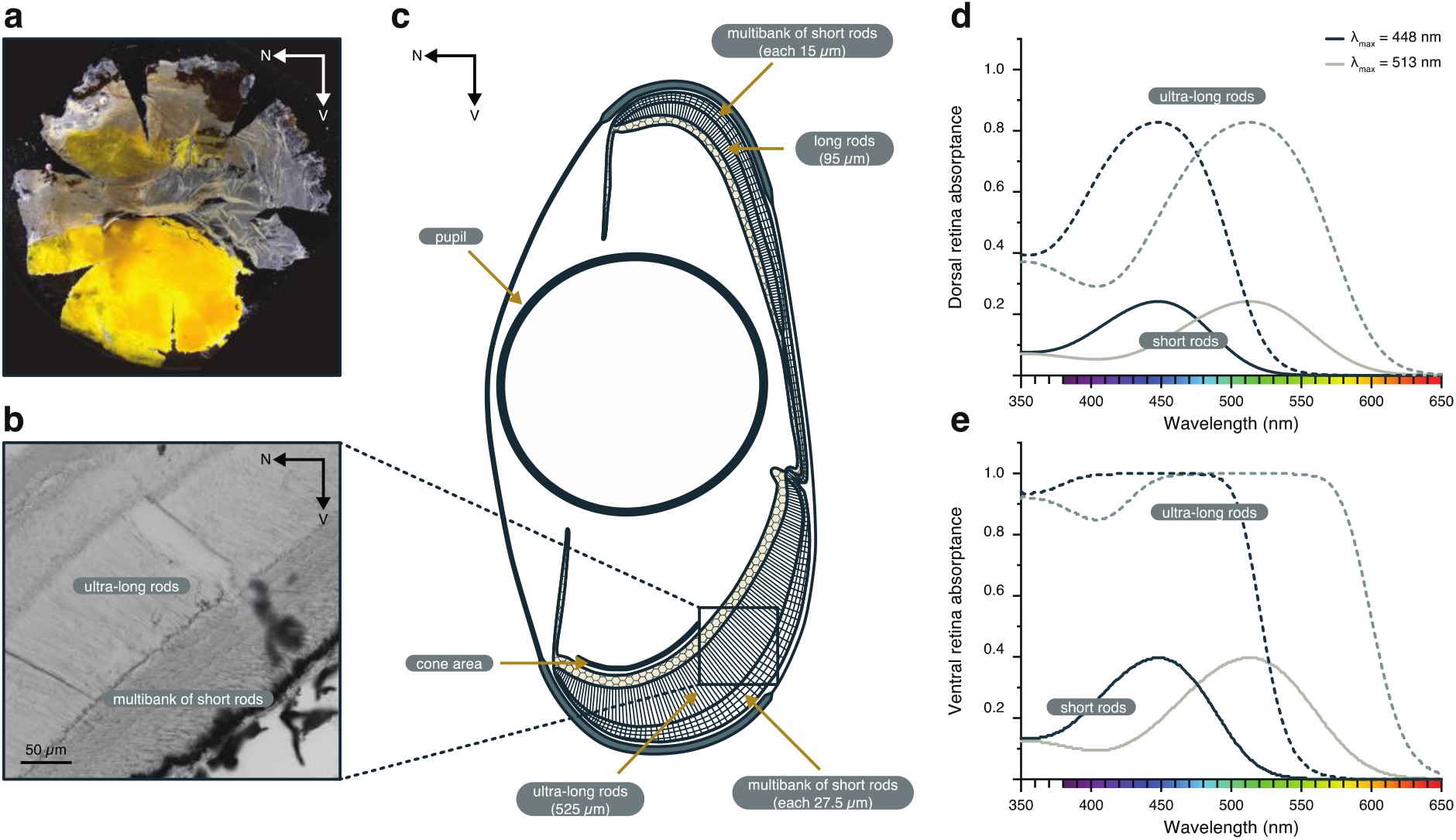
The visual system of the silver spinyfin (*Diretmus argenteus*). **a,** Retinal wholemount depicts distinct differences between the ventral (dark yellow pigment), central (no pigment), and the dorsal (light yellow pigment) areas of the *D. argenteus* retina. **b,** Micrograph showing the ultra-long and the short-rod banks (seven in this case) in the ventral region of the retina. **c,** Schematic of a cross-section of the *D. argenteus* eye adapted from ref. 19. At least six distinct areas can be depicted: ventral and dorsal long and short rods, central area of short rods, ventral area of cone photoreceptors. **d-e,** Modelled photon absorptance of the (ultra) long and short rods in the dorsal (**d**) and ventral (**e**) retina if assuming the single expression of either the shortest or the longest tuned of the *D. argenteus* rod photopigments. Note, that the shortest and the longest tuned photopigment expressed in the ultra-long photoreceptors of the ventral area would be enough to cover photon absorptance across the available ambient light spectrum (in colour).

Second, spectrally distinct RH1 photopigments may be used for colour recognition of some form^20^. Invertebrates are well documented to use differently tuned photoreceptors to enable colour discrimination in dim-light environments^32,33^. For vertebrates, only geckos and frogs have been shown to achieve a similar feat^34–36^. In these cases rod shaped photoreceptors rely on differently tuned cone visual pigments (e.g., in geckos^34^) or a combination of cone pigments and a single rod photopigment (e.g., in frogs^35,36^) to discriminate between colours. Considering the relative simplicity of teleost brains, however, it is unlikely that *D. argenteus* is using colour opponency to compare between 14 differently tuned spectral channels. Instead, a different form of colour perception might be at play. Stomatopod crustaceans, for example, which have a similar number of differently tuned photopigments to the Diretmidae, use scanning eye movements instead of colour opponency to quickly recognize rather than distinguish colours^37^. Although fishes are not known to have such eye movements, the visual scene encountered in the deep-sea is unlike anything else found on earth; deep-sea fishes encounter colour signals in the form of narrow-banded bioluminescent emissions surrounded by darkness^38^. Hence, colour signals are unlikely to be seen in parallel, and quickly recognizing a specific bioluminescent colour might be beneficial for survival.

Supporting the existence of purely rod based colour vision systems in deep-sea fishes, members from all the deep-sea lineages with expanded *RH1* gene repertoires are known to have rod photoreceptors with different spectral sensitivities and additional yellow spectral filters within the dioptric or retinal regions that have been suggested to enhance colour discrimination (Figure 4a)^39,40^. Denton and Lockett (1989)^20^ have even suggested colour vision was possible in the Diretmidae but at that stage identified a single rod visual pigment situated in each of the multibank retinal regions (i.e. one pigment in the dorsal and another one in the ventral retina). A likely driver for colour sense in the deep sea are spectrally diverse bioluminescent spectra^13^. Alternatively, a spectrally optimised single-colour object contrast system may help break bioluminescent camouflage against downwelling light for identifying prey or conspecifics^20^. In a fascinating parallel in the cephalopods, the otherwise expected monochromatic abraliopsid squids also express multiple visual pigment types in multibank retinae and utilise differently coloured bioluminescent signals during seasonal mating^41,42^. By all means, our findings shift the current paradigm of vertebrate vision in terms of the role of rod photoreceptors, irrespective of the exact function these multiple rod opsins have in deep-sea fishes.

## Acknowledgments

We thank Andrew Bentley, Michael Berenbrink, Wen-Sung Chung, Adrian Indermaur, Xabier Irigoien, Stein Kaartvedt, Lukáš Kalous, M. Danielle MacDonald, Jiří Peterka, Genevieve Phillips, Jan Yde Poulsen, Even Sannes Riiser, Anders Rostad, Olivia Roth, Ana Gro Salvanes, and Lee Frey (HBOI/Blue Turtle Engineering) for their help in the field and/or for providing samples; the staff of the Lizard Island Research Station for providing logistical support; the captains and crews of the research vessels *Seward Johnson*, *Walther Herwig III*, *Sonne*, *G. O. Sars*, *Thuwal*, *Maria S. Merian,* and *Trygve Braarud* for providing the opportunity to participate and collect samples on various cruises; Janette Edson, Christian Beisel, and Craig Michell for help with sequencing at the genomics facilities of the Queensland Brain Institute, University of Basel, and KAUST, respectively; the Waitt Foundation for Discovery for hosting and financial support; and Francesco Santini for discussions regarding fossil calibrations.

## Author Contributions

Z.M., F.C., J.M., and W.S. conceived the study. Mi.M., Ma.M., K.S.J., and S.J. performed the genome sequencing. Z.M., F.C., F.d.B., J.M., and W.S. carried out and supervised the transcriptome sequencing. Z.M. and F.C. carried out the opsin gene mining, analyses, and raw-reads assemblies. F.C. and S.M.S. performed the PGLS analyses. Mi.M., Ma.M., O.K.T., and S.J., carried out and supervised the genome assembly. Mi.M. carried out phylogenetic analyses. K.L.C. performed the regulatory region analyses. W.I.L.D. and J.K.M. carried out the *in vitro* protein regeneration and spectral prediction analyses. R.H. provided fish samples for the study. All authors have read and contributed to the final version of the manuscript.

## Data and materials availability

New draft genome sequences of deep-sea fishes are available at https://hts-nonsecure.web.sigma2.no/zuza_fabio/ (and have been submitted to the European Nucleotide Archive, ENA); transcriptomic data that support the findings of this study have been deposited on GenBank (BioProject ID PRJNA421052). Source files including custom scripts and sequence alignments have been deposited in GitHub (https://github.com/mmatschiner/opsin_evolution), phylogenetic trees are available at http://evoinformatics.eu/opsin_evolution.htm. All other data analysed during this study are included as Supplementary Materials. Correspondence and requests for materials should be addressed to, or.

## Supplementary Materials

Material and Methods

Supplementary Text - Extended Material and Methods

Supplementary Text - Ethical approval and relevant permits

Supplementary Text - Fossil calibrations for time calibrating the teleost species tree Figures S1 – S5

Tables S1 – S9

References (43 – 232)

